# To make or take: bacterial lipid homeostasis during infection

**DOI:** 10.1101/2021.01.06.425669

**Authors:** Felise G. Adams, Claudia Trappetti, Jack K. Waters, Maoge Zang, Erin B. Brazel, James C. Paton, Marten F. Snel, Bart A. Eijkelkamp

## Abstract

Bacterial fatty acids are critical components of the cellular membrane. A shift in environmental conditions or in the bacterium’s lifestyle may result in the requirement for a distinct pool of fatty acids with unique biophysical properties. This can be achieved by the modification of existing fatty acids or via *de novo* synthesis. Furthermore, bacteria have evolved efficient means to acquire these energy-rich molecules from their environment. However, the balance between *de novo* fatty acid synthesis and exogenous acquisition during pathogenesis is poorly understood. Here we studied the mouse fatty acid landscape prior and post infection with *Acinetobacter baumannii*, a Gram-negative, opportunistic human pathogen. The lipid fluxes observed following infection revealed fatty acid- and niche-specific changes. Lipidomic profiling of *A. baumannii* isolated from the pleural cavity of mice identified novel *A. baumannii* membrane phospholipid species and an overall increased abundance of unsaturated fatty acid species. Importantly, we found that *A. baumannii* relies largely upon fatty acid acquisition in all but one of the studied niches, the blood, where the pathogen biosynthesises its own fatty acids. This work is the first to reveal the significance of balancing the making and taking of fatty acids in a Gram-negative bacterium during infection, which provides new insights into the validity of targeting fatty acid synthesis as a treatment strategy.

**Importance:** *Acinetobacter baumannii* is one of the world’s most problematic superbugs, and is associated with significance morbidity and mortally in the hospital environment. The critical need for new antimicrobial strategies is recognised, but our understanding of its behaviour and adaptation to a changing environment during infection is limited. Here, we investigated the role of fatty acids at the host-pathogen interface using a mouse model of disease. We provide comprehensive insights into the bacterial membrane composition when they colonise the pleural cavity. Further, we show that *A. baumannii* heavily relies upon making its fatty acids when residing in the blood, whereas the bacterium favours fatty acid acquisition in most other host niches. Our new knowledge aids in understanding the importance of host fatty acids in infectious diseases. Further, fatty acid synthesis is an attractive target for the development of new antimicrobial strategies, but our work emphasizes the critical need to understand the microbial lipid homeostasis before this can be deemed suitable.

## Observation

The bacterial type II fatty acid synthesis (FASII) pathway has been interrogated as a potential antimicrobial target, with the validity of this strategy debated through study of the Gram-positive bacterium, *Staphylococcus aureus* (1, 2). In addition to FASII, pathogens can readily acquire fatty acids from their environment through either the FakAB system (Gram-positive bacteria) (3) or FadL (Gram-negative bacteria) (4, 5), which may render them resistant to FASII-targeting antimicrobial strategies, as illustrated in *S. aureus* (6, 7). Nevertheless, *S. aureus* fatty acid auxotrophs are avirulent in a mouse model of disease (8), indicating that at least some level of *de novo* synthesis is critical to bacterial survival during infection. Interestingly, the balance between *de novo* fatty acid synthesis and exogenous acquisition has never before been studied in Gram-negative bacterial pathogens. Further, the niche-specificity of bacterial lipid homeostasis is poorly understood.

We defined the fatty acid composition of various mouse niches and their changes following intranasal challenge with the Gram-negative human pathogen *Acinetobacter baumannii* (strain AB5075_UW) (**Supplementary Methods**). Overall, the fatty acid abundance in mouse plasma decreased following infection, with the most dramatic changes seen in 16:0, 18:1 and 18:2 (**Fig 1a**). In contrast, 20:4 was not affected by *A. baumannii* infection, potentially due to a role in immune-activation; findings consistent with those in mice infected with *Streptococcus pneumoniae* (9). The liver is the primary site of host fatty acid synthesis and its profiling revealed the most pronounced increases in 18:1 and 18:2 (**Fig 1b**), possibly to replenish their deficit observed in plasma. The bronchoalveolar lavage (BAL) represents the primary bacterial challenge site, where 16:0 is the predominant fatty acid (**Fig 1c**). The 16:0 abundance decreased marginally following infection, whereas that of 18:1 increased. Finally, we assayed the pleural lavage (PL), a sterile niche in uninfected animals with fatty acid levels below detection (**Fig 1d**). *A. baumannii* colonisation resulted in the subsequent identification of 16:0, 18:2, 18:0, 18:1 and 20:4 species (in order of abundance). Similar to that observed in mouse plasma, the increase in 20:4 in the PL may be due to a contribution to immune modulation. Overall, these findings indicate that *A. baumannii* necessitates adaptation to vastly different lipid landscapes when colonising distinct host niches. The changes in the fatty acid profiles in these niches following infection may be a result of the host response to infection, in combination with the bacterium acquiring and secreting fatty acids during proliferation.

**Figure 1.**
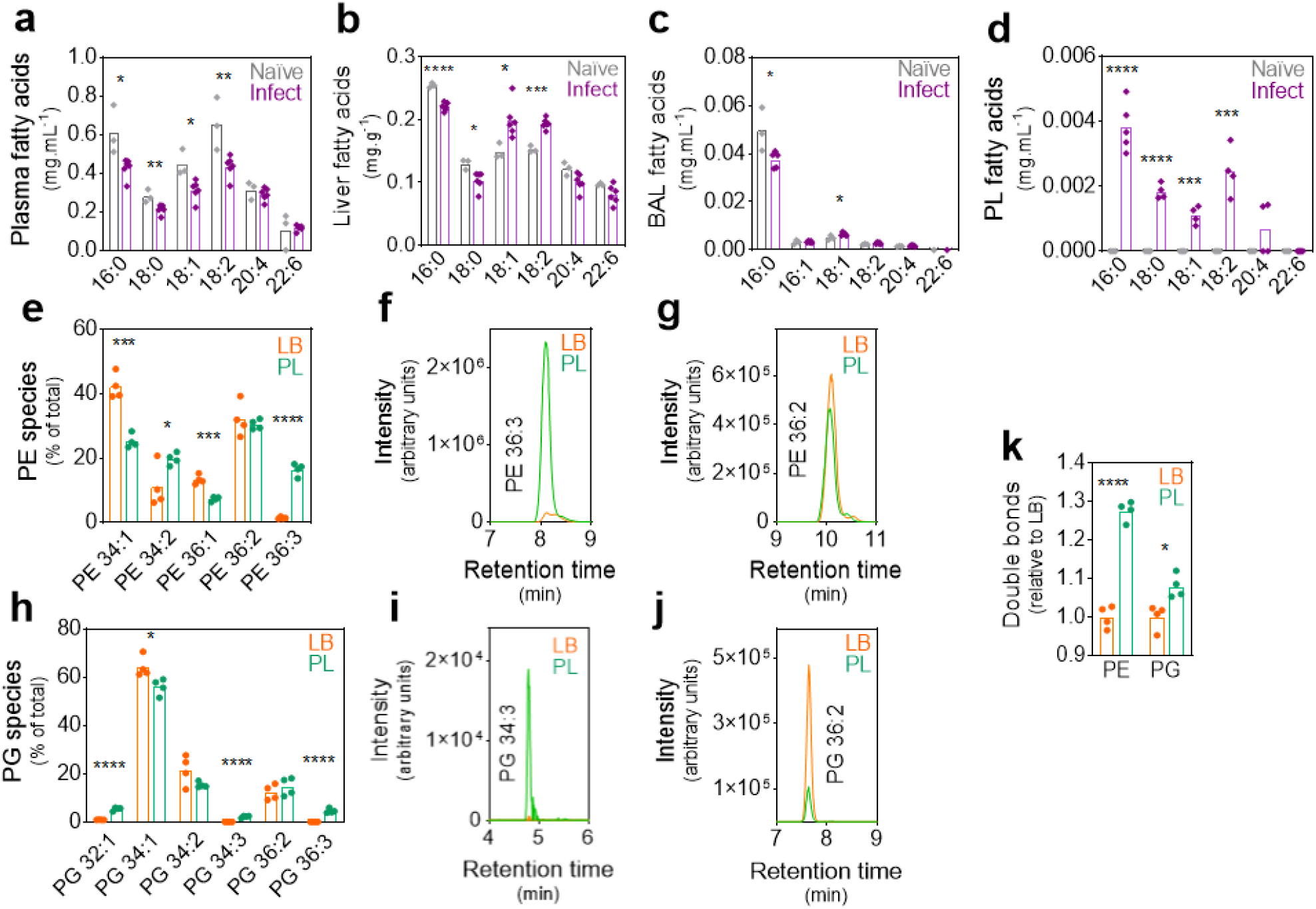
The host and *A. baumannii* lipid landscape. The fatty acids (milligram of fatty acid per millilitre or gram of tissue) in the plasma (**a**), liver tissue (**b**), bronchoalveolar lavage (BAL) (**c**) and pleural lavage (PL) (**d**) of 9-week old female BALB/c mice were examined prior (Naïve; grey) and 24 hours post intranasal challenge with *A. baumannii* strain AB5075_UW (Infect; purple). The phosphatidylethanolamine (PE) (**e**) and phosphatidylglycerol (PG) (**h**) species (number carbons: number of double bonds in the acyl chains) were quantified in *A. baumannii* cultured in Luria-Bertani media (LB) or *A. baumannii* from the pleural lavage of BALB/c mice 24 hours post intranasal challenge (PL), using liquid chromatography-mass spectrometry (LC-MS). The LC-MS chromatograms for heavily enriched lipid species in *A. baumannii* isolated from the PL [PE 36:3 (**f**) and PG 34:3 (**i**)], and control lipid species present in *A. baumannii* from the PL and those cultured in LB [PE 36:2 (**g**), PG 36:2 (**j**)]. The data represent the average of 4 biological replicates. The relative number of double bonds of PE and PG species in *A. baumannii* from LB or the PL was defined using the percentage abundance (**k**). All statistical analyses were performed using Student’s *t*-tests; * p ≤ 0.05, ** p ≤ 0.01, *** p ≤ 0.001 and **** p ≤ 0.0001.

To ascertain the potential exchange of fatty acids between *A. baumannii* and the host environment, we analysed the lipidome of *A. baumannii* isolated from the pleural cavity and compared this to *A. baumannii* cultured in Luria-Bertani media (LB) (**Supplementary Methods**). Unlike the BAL, the PL is sterile in uninfected animals, which eliminates contamination of the sample by non-*A. baumannii* bacterial species. Further, as *A. baumannii* colonisation in this niche is high (>10^9^ CFU per sample), there is no need for tissue disruption or removal of erythrocytes and host immune cells can easily be depleted from the lavage via centrifugation. We studied the most abundant *A. baumannii* phospholipids, phosphatidylethanolamine (PE) and phosphatidylglycerol (PG), by liquid-chromatography-mass spectrometry (LC-MS). The most striking differences were observed in PE 34:1 (1.7-fold decrease) and PE 36:3 (11-fold increase) when compared to *A. baumannii* grown in LB media (**Fig 1e,f**). Similar to PE 36:3, which consists of 18:1 and 18:2 acyl chains, PG 34:3 and PG 36:3 contain 18:2 acyl chains and were identified in *A. baumannii* from the PL but not in LB (**Fig 1h,i**). PG 34:3 (16:1 + 18:2) is an unusual species, not commonly found in mammalian cells (LipidMAPS database), nor in *A. baumannii* grown in LB media (**Fig 1i**) (10). Hence, its presence in *A. baumannii* isolated from the mouse pleural cavity is a strong indicator of *A. baumannii* 18:2 acquisition from the host, reminiscent of 18:1 acquisition by *S. aureus* in mouse thigh tissue (11). The enhanced incorporation of 18:2 fatty acids in the *A. baumannii* PE and PG pools affected the total number of unsaturated fatty acids in *A. baumannii* from the PL, as compared to those cultured in LB, by 1.3-fold (PE) and 1.1-fold (PG) (**Fig 1k**).

To study the relative contribution of fatty acid acquisition to *A. baumannii* virulence, we challenged mice intranasally with AB5075_UW wild-type and a *fadL* mutant derivative (*A. baumannii fadL*::T26) and compared their invasion in diverse niches. The *fadL* mutant was significantly impaired in the colonisation of the bronchoalveolar lumen, lung tissue, pleural cavity, liver and spleen, ranging from 6.5 × 10^2^-fold (BAL) to 2.0 × 10^4^-fold (PL) (**Fig 2a**). In contrast, colonisation of the *fadL* mutant was not significantly different to the wild-type in the blood (1.6 × 10^1^-fold, *p* = 0.101). Although these phenotypes are most likely related to the role in fatty acid acquisition, FadL is a surface-exposed protein that has also been shown to play a role in adherence to host cells in *Haemophilus influenzae* (12). Hence, from these data alone we cannot rule out a multi-factorial cause behind the observed colonisation differences.

**Figure 2.**
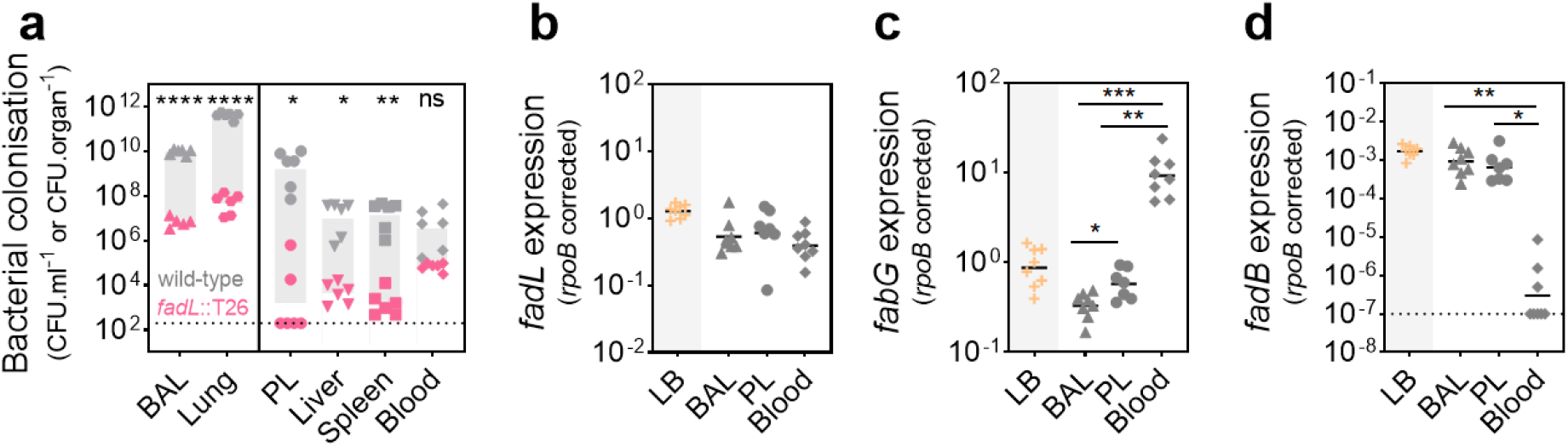
The niche-specific balance between fatty acid acquisition and *de novo* synthesis. (**a**) The role of FadL-mediated fatty acid acquisition in *A. baumannii* colonisation was examined following intranasal challenge of 9-week old female BALB/c mice with 2 × 10^8^ colony forming units of *A. baumannii* AB5075_UW (wild-type; grey) and the *fadL* mutant (*fadL*::T26; pink). The difference between the two groups (geometric means) is indicated with the grey bar. The gene expression of *fadL* (**b**), *fabG* (**c**) and *fadB* (**d**), corrected against those defined for *rpoB*, in *A. baumannii* AB5075_UW cultured in Luria-Bertani media (LB) or isolated from the bronchoalveolar lavage (BAL), pleural lavage (PL) or blood, was examined by qRT-PCR. For all panels, the dotted line indicates the limited of detection (**a** and **d**). All statistical analyses were performed using Student’s *t*-tests; * p ≤ 0.05, ** p ≤ 0.01, *** p ≤ 0.001 and **** p ≤ 0.0001; ns = not significant.

To delineate the role of exogenous fatty acid acquisition relative to *de novo* synthesis, we examined the transcription levels of *fadL* and *fabG* (a critical FASII member) in *A. baumannii* from the BAL, PL and blood (**Supplementary Methods**). Overall, *fadL* transcription was similar in distinct host niches (**Fig 2b**). Although, *fabG* expression was significantly higher in *A. baumannii* from the PL as compared to the BAL (1.8-fold), *fabG* transcription was 17-fold greater in *A. baumannii* from the blood compared to the PL (**Fig 2c**). In combination with FadL being near-superfluous during *A. baumannii* bacteraemia, this is a strong indicator that *A. baumannii* relies heavily upon *de novo* fatty acid synthesis whilst residing in the blood. In other niches, *A. baumannii* is likely to augment the intracellular pool of fatty acids with exogenously acquired fatty acids through uptake via FadL. To gain greater insights into the intracellular *A. baumannii* lipid status we also assayed the expression of *fadB* (a critical component of the β-oxidation pathway), as its expression in *A. baumannii* is associated with an intracellular fatty acid surplus (10). Interestingly, whereas *fadB* expression in *A. baumannii* was comparable in the PL and BAL, in the blood, expression decreased by approximately 700-fold (**Fig 2d**). This is consistent with *A. baumannii* experiencing fatty acid limitation and reliance upon synthesising its own fatty acids in this niche. Transcriptional profiling of *fadL*, *fabG* and *fadB* in *A. baumannii* cultured in LB indicated that the bacterium balances *de novo* synthesis and fatty acid uptake/recycling when grown in this medium as it was analogous to that observed in *A. baumannii* in the PL (**Fig 2b-d**).

## Conclusions

*A. baumannii* is an opportunistic pathogen, with limited host-adaptation traits. Here we have illustrated that *A. baumannii* utilises environmental fatty acids to promote colonisation in various niches of the host, which includes non-self fatty acids, such as 18:2, to generate a unique phospholipid profile. Host fatty acid profiling revealed a decrease in most fatty acid species in the plasma following infection with *A. baumannii*. This could be a host-mediated response to infection or a depletion due to *A. baumannii* acquisition during early stages of bacteraemia. However, the relative competitiveness of the *fadL* mutant compared to the wild-type in the blood suggests the former. Despite the relatively high abundance of fatty acids in the plasma, these do not appear to be bioavailable to *A. baumannii*, potentially as a result of sequestration by plasma lipid binding proteins such as albumin and low/high-density lipoproteins. Fatty acid synthesis is an attractive target for the development of new antimicrobial strategies, and our work has underscored the critical need to understand microbial lipid homeostasis before this can be deemed suitable. Overall, FASII-targeting antibiotics are likely to be most effective against *A. baumannii* bloodstream infection.

## Acknowledgements

This work was supported by the National Health and Medical Research Council (Australia) through Project Grant 1159752 to BAE. MZ is a recipient of an Australian Government Research Training Program scholarship. We are grateful to Jake White (South Australian Health and Medical Research Institute) for his technical assistance with the phospholipid analyses and discussions. We thank Robert Gibson and Kristina Hickson (South Australian Health and Medical Research Institute) for their input into the host fatty acid analyses.

## Supplementary Material

Supplementary methods

